# Evaluating Molecular Docking Programs for RNA-Targeted Ligand Screening: Influence of Binding Modes and Ligand Types

**DOI:** 10.1101/2025.07.03.661502

**Authors:** Kaichen Wang, Louis DeFalco, Chaitanya K. Jaladanki, Roland G. Huber, Christina Li Lin Chai, Hao Fan

## Abstract

Structured RNAs are critical regulators of gene expression and represent promising targets for drug discovery. Despite this potential, structure-based virtual screening approaches—in particular, molecular docking—have been underutilized for RNA targets, largely due to the intrinsic conformational flexibility of RNA and the limited docking methods specifically optimized for RNA. In our study, we evaluated the efficacy of three widely used docking programs for RNA-ligand docking screens. The performance of these programs — Glide, Gold, and rDock, were evaluated with the dataset of 25 diverse RNA targets with 432 small molecule RNA binders. We found that the performance was similar across docking methods, but consistent differences emerged among RNA classes. We further explored consensus scoring across both receptor conformations and docking methods, finding that selecting the average scores across docking programs and RNA structures yielded stable performance when compared to individual methods. Finally, clustering analysis of RNA binders revealed distinct scoring function preferences for different chemotypes of RNA binders, underscoring the potential for tailored scoring strategies in RNA-targeted drug discovery. Together, this work provides a comprehensive evaluation of the performance of available docking methods in different cases of RNA-ligand docking and offers practical guidance for improving virtual screening against structured RNAs.

## INTRODUCTION

RNA plays essential roles in gene expression processes, including scaffolding, catalysis and regulatory roles in gene translation, making it an attractive target for drug discovery. Despite its limited chemical diversity with only four nucleosides, RNA can adopt intricate secondary and tertiary structures, presenting both opportunities and challenges for structure-based drug design. In addition to emerging RNA-targeting strategies such as CRISPR (1), antisense oligonucleotides (2), and small interfering RNAs (3) that focus on large biomolecules, small molecules can also bind to structured RNA and modulate their biological functions. For instance, several classes of first-line antibiotics, including aminoglycosides, macrolides and tetracyclines, target structured regions in bacterial ribosomal RNA to inhibit gene translation (4). As regulatory structured RNA elements commonly identified in bacterial genomes, riboswitches can modulate the expression of specific genes through the change of tertiary structure caused by the binding of particular metabolites (5,6). In human cells, structured RNAs, such as long noncoding RNA and microRNA, play key roles in mRNA splicing, cell proliferation and progression of cancer, and have been identified as targets of small molecules. One such example is Malat1, a long noncoding RNA element that regulates the progression of lung cancer, which was selectively targeted by small molecules and inhibited lung cancer progression in mice (7). Overall, RNA-targeting small molecules have shown significant potential to modulate RNA function, providing an alternative strategy for drug discovery, particularly for targets deemed undruggable on the protein level.

Among these emerging cases of small molecules that target structured RNAs, structure-based computational methods have been applied to RNA targets. Molecular docking and pharmacophore-based virtual screening have been used to identify novel binders to guanine riboswitch (8), preQ1 riboswitch (9), HCV IRES (10), TPP riboswitch and HIV-TAR (11–13). In another study by Ganser et al. (14), the intrinsic flexibility of structured RNA was taken into account, and an ensemble of HIV-TAR conformations was generated using molecular dynamics simulations. Subsequent screening of a compound library against this ensemble of conformations led to the discovery of ligands with novel scaffolds. However, despite these advances, structure-based computational methods have not been as extensively applied as experimental approaches, such as high-throughput screening, for identifying small molecule ligands targeting RNA.

Several issues restrict the further application of structure-based computational techniques to RNA. RNA exhibits high structural flexibility and, like the induced-fit model of enzymes, undergoes broad conformational changes upon ligand binding (15). Additionally, the negatively charged phosphate RNA backbone interacts frequently with metal ions, which are essential for structural stabilization and, in some cases, mediate RNA-ligand interactions (16). Despite these challenges, computational methods are increasingly being optimized for RNA over time. Docking programs that were primarily developed for protein-ligand interactions, such as Glide (17), GOLD (18), DOCK (19) and ICM (20), have demonstrated promising results in RNA-ligand docking, despite not being specifically optimized for RNA targets. RiboDock (21) and its successor rDock (22) were among the first docking tools that were optimized against both protein and nucleic acid targets, with rDock offering refined scoring schemes. To complement the available docking programs, scoring functions specifically tuned for RNA-ligand interactions such as SPA-LN (23), ITScore-NL (24), SPRank (25) and DrugScoreRNA (26) were developed. Besides molecular docking, other standalone methods including RLDock (27), RNAAmigos (28), RNAPosers (29), ANNAPURNA (30), integrate empirical and machine learning approaches to predict RNA-ligand binding. Physics-based techniques, including molecular dynamics (MD) simulations, free energy perturbation, and MM-GBSA/PBSA, further refine RNA-ligand complexes (31–33). The growing availability of nucleic acid structure databases (HARIBOSS (34), NAKB (35)) and RNA binders (R-BIND (36), R-SIM (37), NALDB(38)) provide essential datasets of structures and ligands to further support RNA drug discovery efforts.

With the advancement of RNA-targeted small molecule drug discovery, accurately evaluating the performance of docking methodologies has become increasingly crucial. While previous studies, such as the ones by Kallert (9), Gunasinghe (39), Li (40), and Chen (41) have assessed docking methods for RNA-ligand docking, there are no studies that benchmark on a comprehensive dataset with the scale and organization comparable to established databases. To address this gap, we compiled a dataset of 226 ligand-bound RNA structures from the Protein Data Bank (www.rcsb.org) (42) and 432 experimentally validated RNA binders, covering 25 RNA targets (Figure 1). The RNA structures and binders were classified based on sequence clustering of each target RNA. Based on the annotation of the NAKB database (35,43) that describe the RNA-ligand interactions, 24 of the 25 RNA targets were further grouped by ligand binding modes into the following: groove binding (ligands occupying the RNA grooves), intercalating (ligands inserting between base pairs), and nucleotide-like binding (ligands mimicking natural nucleotides via stacking and hydrogen bonding in the helix). Using this dataset, we docked the 432 RNA binders and corresponding DUD-E (44) decoys against 25 PDB structures with three widely used docking programs: Glide, Gold, and rDock. Docking performance was evaluated in terms of pose prediction and the enrichment of RNA binders over decoys. Additionally, we compared the performance across ligands of different chemotypes and RNA target types to identify factors that influence docking performance. Altogether, our benchmarking study reveals that while Glide, GOLD, and rDock achieve similar average enrichment across 25 RNA targets, their relative performance varies significantly for different binding types, with docking against RNA targets associated with intercalation and nucleotide-like binding performing consistently better than those associated with groove binding. We also demonstrate that consensus scoring yields stable early enrichment as compared to individual methods, particularly averaging across programs. Furthermore, the observed variation in performance across ligand chemotypes suggests that specific scoring functions may be better suited to particular chemical classes. This comprehensive benchmarking study provides practical guidance for optimizing virtual screening workflows to evaluate chemically diverse libraries against a broad range of structured RNA targets in future studies.

**Figure 1.**
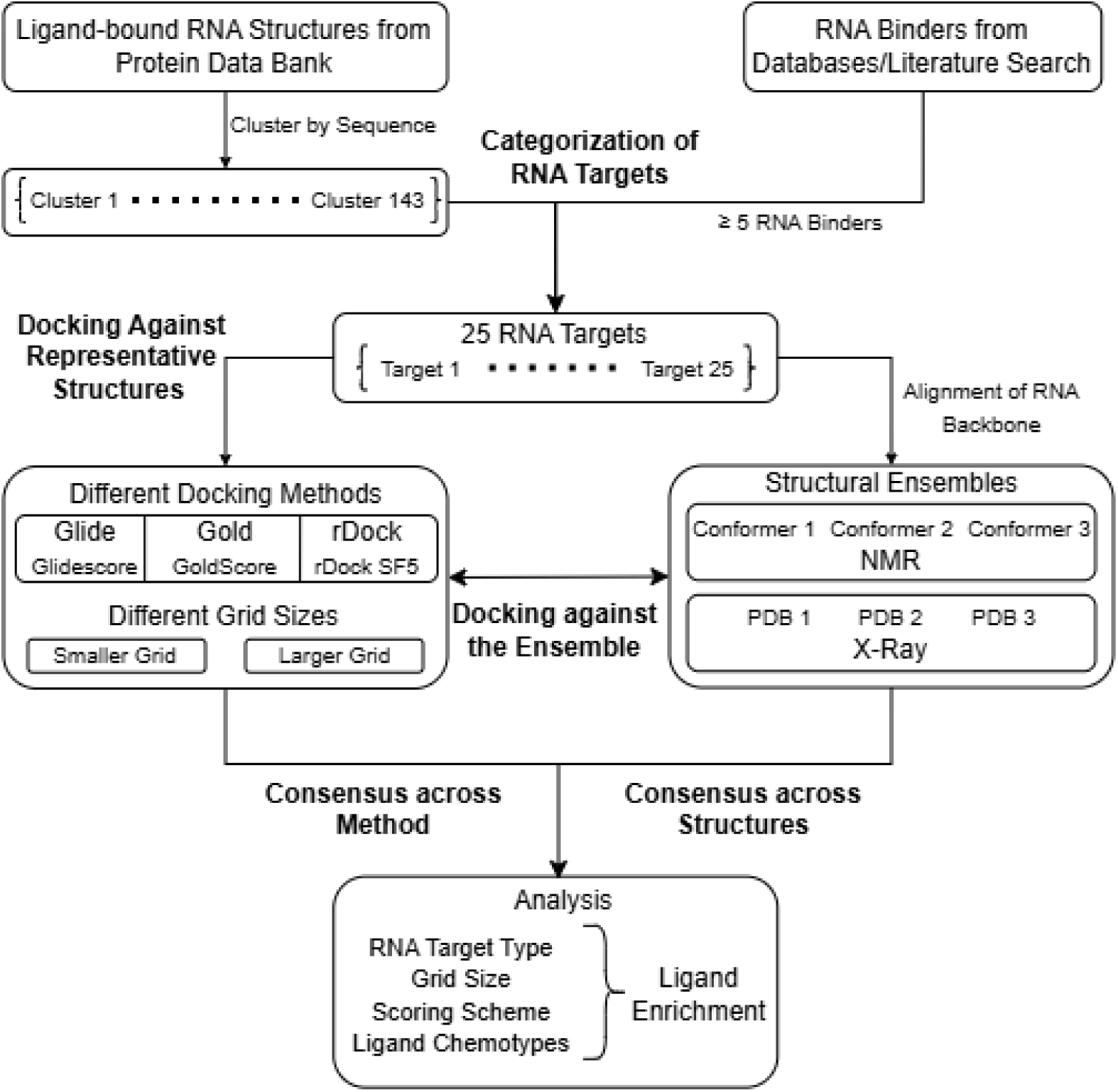
Workflow of benchmarking molecular docking methods for RNA-ligand docking. A dataset of 25 structured RNA targets was curated through literature search and sequence-based clustering. RNA binders were docked against one representative RNA structure for each target. For six selected targets, two additional conformers were included, chosen based on RNA backbone alignment to assess conformational variability. Docking performance was primarily evaluated by ligand enrichment, and trends were analyzed with respect to RNA target type, docking grid size, scoring scheme, and ligand chemotype.

## MATERIAL AND METHODS

### RNA Dataset Generation

A customized, curated database of small molecule ligand-bound RNA structures was retrieved using publicly available biomolecule structure databases and from literature sources. 3797 PDB structures containing keywords “RNA” and “ligand” were downloaded from the RCSB Protein Data Bank (https://www.rcsb.org/). Structures that were complex with only ions and/or crystallization agents were removed to isolate unique RNA-ligand complexes from the dataset, reducing the size of the dataset by approximately 50%. Then, the RNA-protein complexes with ligands bound to RNA were kept, and the remaining ones were removed from the dataset, further reducing the dataset to less than 20% of the original entries. Furthermore, all the DNA/RNA hybrid structures and large ribosomal protein-RNA complexes were removed, yielding 538 unique RNA-ligand complexes. The RNA sequences of the curated 538 ligand-bound RNA structures were extracted from the .cif files using the Biopython 1.83 MMCIF2Dict module (45). RNA sequences were then clustered with CD-Hit (46) using a similarity cutoff of 85%, which gave 143 clusters (Table S1).

The RNA binders were collected from dedicated databases such as R-Bind, R-Sim, and HARIBOSS, and literature search (9). Each RNA binder must have information regarding its target RNA sequence and have binding affinity data (e.g., K_d_, IC_50_, EC_50_, etc.). The RNA binders without binding affinity data or target RNA sequence were removed. Only ligands with affinity ≤100µM were included in the dataset. The RNA binders with molar weight higher than 1000 Da (e.g., adenosylcobalamin) or lower than 100 Da (e.g., guanidine) or having less than 5 heavy atoms (e.g., metal ions) were not considered. The list of RNA binders is included in Table S2, along with the physicochemical properties generated by Maestro (47), SwissAdmet (48) and the Python package rdKit (2023.3.3).

### Classification of RNA Targets

The PDB structures under each of the 143 clusters were sorted into RNA targets based on the description. For example, all 16 PDB structures in cluster 19 were labelled as “FMN riboswitch” based on their PDB titles. Thereafter, the target sequence of each RNA binder was aligned and matched against the sequences of ligand-bound RNA PDB structures using JalView (49). This process identified 25 RNA targets across different clusters, each containing at least five RNA binders. Notably, the list contains RNA targets that are closely associated but have different sequences, namely: THF-I and THF-II riboswitches, SAM-I and SAM-SAH riboswitches, eukaryotic 16s rRNA A-site and prokaryotic 16s rRNA A-site.

The 25 RNA targets were classified based on the patterns of RNA-ligand interactions according to the annotations on the NAKB database: intercalating, nucleotide-like binding, groove binding, and miscellaneous. An alternative classification of RNA targets was also applied based on their biological functions under 5 main categories: artificial aptamers, disease-related human RNA targets, bacterial ribosomal RNA targets, bacterial riboswitch, and viral RNA targets (Table 2). For each of the 25 unique sequences, one representative X-Ray PDB structure of the same sequence with the highest resolution was then picked for docking. In the case where the X-Ray structure is not available, the first conformer of the NMR structure with the best clashscore (50) with the same sequence was then chosen.

### Molecular Docking

All ligands and decoys in the benchmarking dataset were prepared using LigPrep (51). RNA structures were processed with the Protein Preparation Wizard (52) under the S-OPLS force field. Solvent molecules, ions and cofactors within 6.0 Å of the ligand were removed.

For docking, two parameter sets with different grid sizes were applied across three programs. For Glide, the default scoring function Glidescore was applied, and the precision was set to standard precision (SP). The smaller and larger grid sizes in the parameter were defined by inner boxes of 10 Å and 22 Å edge lengths, respectively, both centered on the center of mass of the reference ligand. The outer box was fixed at 25 Å for both parameters. GoldScore was used as the scoring function for Gold, with other parameters besides cavity box size kept as default. Cavity definitions were set at 6 Å and 12 Å around the reference ligand for the smaller and larger cavity sizes, respectively. For rDock, SF5 was applied as the scoring function with default settings, and cavity definitions followed the same 6 Å and 12 Å thresholds around the reference ligand. Performance of virtual screening was evaluated using the logarithmic area under the curve of enrichment plot (logAUC), which emphasises the early ligand enrichment (53,54). For each RNA target, to calculate the consensus enrichment across three docking methods, docking scores of each ligand or decoy from three methods were standardized using Z-scores (scikit-learn, Python) (55), and logAUC was calculated using either the best or the average Z-score for each ligand and decoy. For docking against multiple structures, all X-ray structures with the same sequence as the representative structure and all NMR conformers under the same representative structure were selected for each RNA target. For each set of structures, residues within 6 Å of the ligand were aligned in UCSF ChimeraX (56) MatchMaker (Needleman–Wunsch algorithm, Nucleic Acid matrix). Based on the RMSD values from alignment, two additional structures were selected so that the sum of RMSD between each of the three structures was maximized. To calculate the consensus enrichment across different RNA structures, for each combination between RNA target and docking method (6 targets * 3 methods), the logAUC was calculated based on the best or the average score of each ligand or decoy against the three structures.

### Clustering of RNA Binders

A Morgan fingerprints for all 432 RNA binders were generated by Python script using rdKit (2023.3.3), from which a fingerprint Tanimoto similarity matrix between each pair of binders was calculated. The similarity matrix was transformed using the UMAP approach with scikit-learn. Based on the resulting UMAP projection, clustering was performed using the HDBSCAN Python module (57) with the following parameters: minimum cluster size of 20, minimum sample of 4, and clustering selection method is leaf. After clustering, the outliers were merged into the nearest clusters with the Tanimoto distance cutoff of 0.7.

## RESULTS AND DISCUSSION

### Accuracy of Pose Prediction

For each of the 25 RNA targets, we select one representative PDB structure with high quality and suitable for docking. For targets with X-ray PDB structures, the structure with highest resolution was selected, and its native ligand was extracted and redocked to its respective RNA structure. For NMR structures, the first conformer with the lowest clashscore (50) in PDB was utilized for molecular docking. Molecular docking was performed using two different grid sizes, smaller and larger grids (as outlined in the methods section), with the results presented in Table 1 and Table S3, respectively. As shown in Table 1, all three docking programs reliably reproduced binding poses similar to native crystal/NMR structures, with RMSD < 2 Å for 40–48% and 44–56% of structures in smaller and larger grid sizes across the docking programs. Among the three methods, Gold performed best with the larger grid, achieving the lowest mean RMSD (3.42 Å) and with RMSD < 2 Å for 48% of the structures, outperforming Glide (4.43 Å, 40%) and rDock (4.09 Å, 40%). In contrast, rDock was the most accurate with the smaller grid, yielding the lowest mean RMSD (3.02 Å) and with RMSD < 2 Å for 56% of the structures, compared to Glide (52%) and Gold (44%). Across both grid sizes, the smaller grid consistently resulted in lower mean RMSD values: 3.70, 3.44, and 3.02 Å for Glide, Gold, and rDock, respectively, compared to 4.43, 3.42, and 4.09 Å with the larger grid. These results indicate that a smaller grid improves pose prediction accuracy, particularly for rDock.

**Table 1.** Summary of re-docking performance for Glide, Gold and rDock against 25 PDB structures.

|  |  |  |  |
| --- | --- | --- | --- |
| RNA Target | Glide | Gold | rDock |
| **Larger Grid Size** |  |  |  |
| Mean RMSD (Å) | 4.43 ± 3.59 | **3.42 ± 2.8** | 4.09 ± 3.45 |
| Success rate (< 2 Å) | 40.00% | **48.00%** | 40.00% |
| **Smaller Grid Size** |  |  |  |
| Mean RMSD (Å) | 3.70 ± 3.60 | 3.44 ± 2.93 | **3.08 ± 2.89** |
| Success rate (< 2 Å) | 52.00% | 44.00% | **56.00%** |
| The best performing item for each category is shown in bold. |  |  |  |

Besides docking parameters, ligand flexibility also affected the docking pose prediction accuracy. For example, as shown in Figure 2, the selected rDock docking poses for RNA targets under three different categories indicates that with smaller grid size, the pose prediction for ligands with less rotatable bonds (Figure 2. A-C, D-F, G-I, with 2, 0 and 2 rotatable bonds respectively)were less accurate compared to those with less rotatable bonds (Figure S1. A-C, D-F, G-I, with 5, 7 and 19 rotatable bonds respectively). This observation is further supported by the RMSD values across 25 structures: ligands with more than five rotatable bonds are associated with higher average RMSD values than those with fewer than five rotatable bonds. For the smaller grid size, average RMSD for the former and later groups are: Glide, 5.99 Å vs. 1.89 Å; Gold, 4.73 Å vs. 2.42 Å; and rDock, 4.45 Å vs. 1.89 Å. A similar trend was observed for the larger grid size: Glide, 6.31 Å vs. 2.95 Å; Gold, 4.26 Å vs. 2.75 Å; and rDock, 6.23 Å vs. 2.55 Å. Despite the limited sample size, the results suggest that the three methods predict the poses with less accuracy when docking more flexible ligands, which requires more precautions in future RNA-targeting ligand discovery studies.

**Figure 2.**
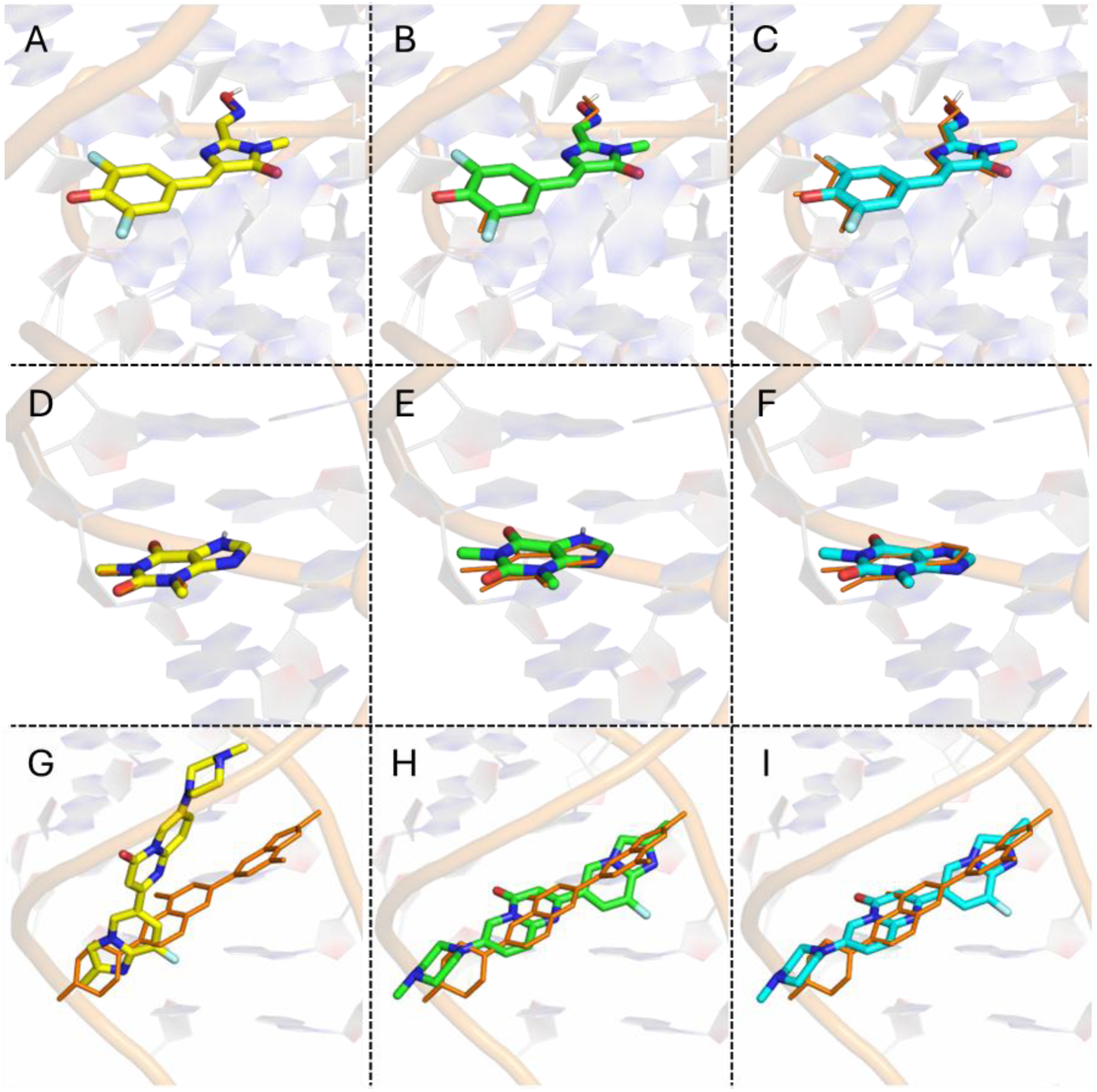
The experimental structures (orange) and binding poses predicted by Glide (yellow), Gold (green) and rDock (cyan) of the native ligands from three classes of RNA targets with different binding modes. (A, B, C) For RNA targets associated with intercalating binding: corn aptamer (PDB: 5BJO). (D, E, F) For RNA targets associated with nucleotide-like binding: theophylline aptamer (PDB: 8D28). (G, H, I) For RNA targets associated with groove binding: SMN2 pre-mRNA (PDB: 6HMO).

### Comparing the Effect of Grid Sizes on Ligand Enrichment

The docking performance is shown in Figure 3 and Table 2 for the smaller grid, and in Figure S2 and Table S4 for the larger grid. Using a 10% difference in logAUC as a practical threshold, ligand enrichment was better with the smaller grid in 3, 7, and 10 cases for Glide, Gold, and rDock, respectively. For example, in the adenine riboswitch, all three methods performed better with the smaller grid, achieving logAUC values of 53.22 (Glide), 33.49 (Gold), and 39.40 (rDock), compared to 38.47, 26.07, and 27.87 with the larger grid. Conversely, the smaller grid performed worse in 4, 7, and 4 targets for Glide, Gold, and rDock, respectively. While for none of the RNA targets the ligand enrichment from the larger grid is better than that from the smaller grid for all three methods, Glide yielded better ligand enrichment against the theophylline aptamer for the larger grid compared to the smaller grid (logAUC 37.04 vs. 43.36). Similarly, Gold performed better with the larger grid for HIV FSS (logAUC of 31.34 vs. 23.50), while rDock performed better with the smaller grid for the THF-II riboswitch (23.40 vs. 27.58).

**Figure 3.**
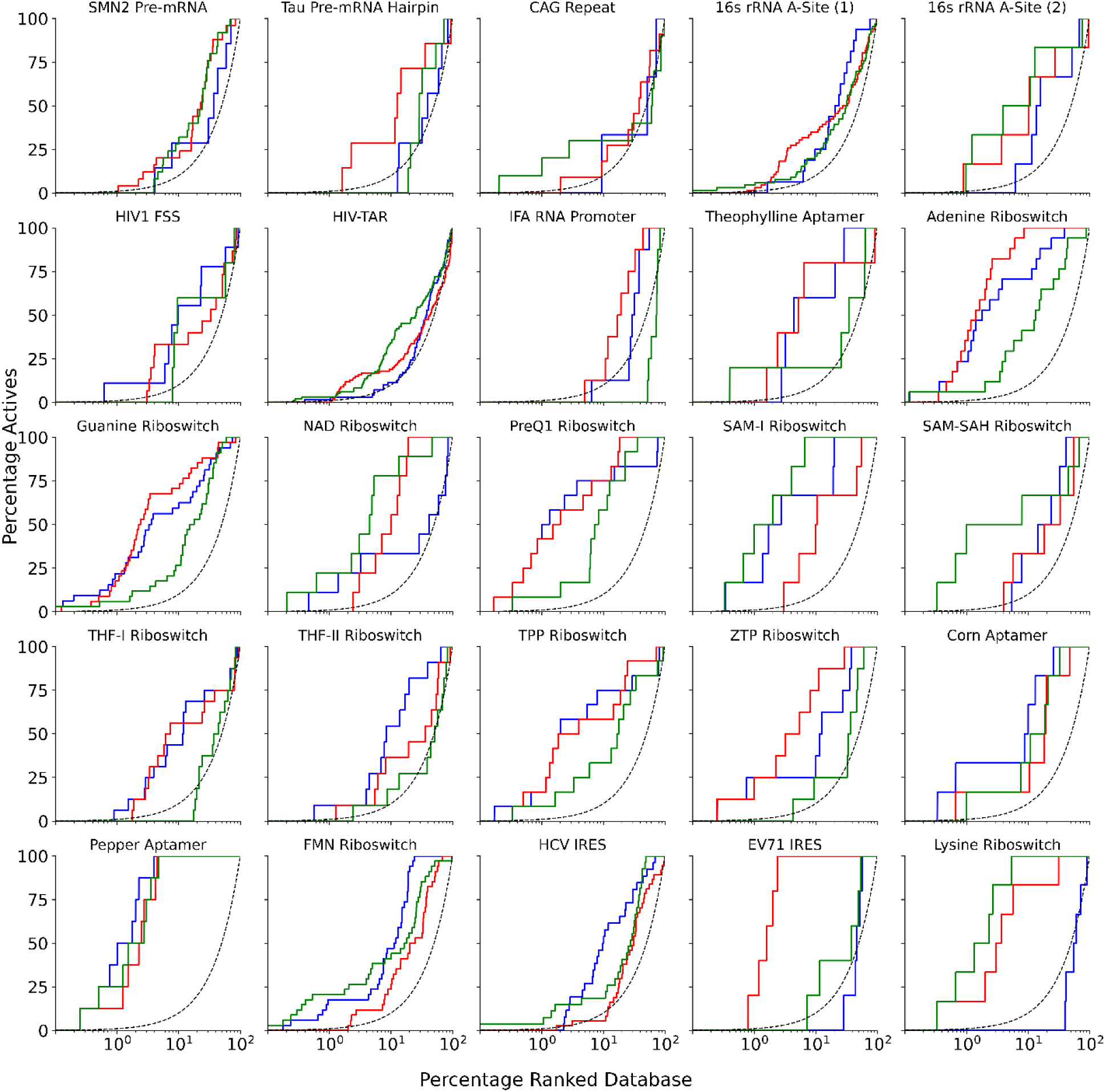
Enrichment curves (logAUC) across the 25 RNA targets using Glide (blue), Gold (green) and rDock (red), with smaller grid size for docking, and random selection (dotted line).The logAUC metric evaluates enrichment by calculating the area under the enrichment curve above the x-axis; curves positioned toward the upper left indicate better early recognition of the RNA binders.

**Figure 4.**
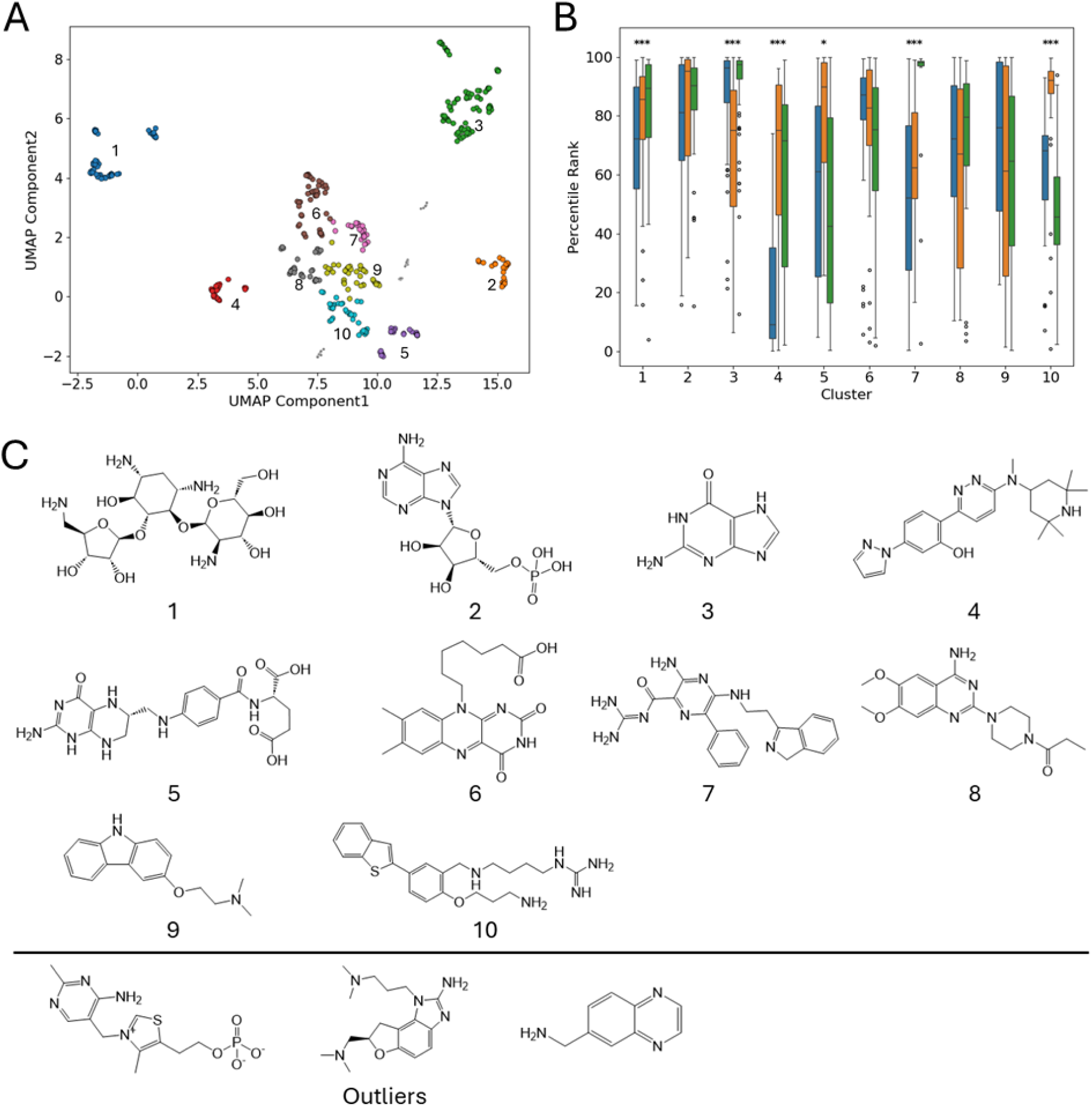
(A) HDBSCAN clustering of RNA binders visualized with uniform manifold approximation and projection (UMAP). The clustering was performed based on Morgan fingerprint similarity matrix. (B) The distribution of percentile ranks from three docking methods for RNA binders from each cluster. The distribution of percentile ranks of RNA binders for Glide, Gold and rDock were shown in blue, orange and green, respectively. Statistical differences between the ranks from different methods were evaluated using Kruskal-Wallis test. * denotes p < 0.05, and *** denotes p < 0.001. (C) The chemotypes of molecules from each cluster and the outliers.

**Table 2.** The docking performance from three individual docking methods and the consensus enrichment based on normalized z-score against 25 RNA targets. Performance is evaluated by logAUC.

| RNA Target | Binding Modes | PDB | Structure | Ligand | Methods |  |  | Consensus Enrichment |  |
| --- | --- | --- | --- | --- | --- | --- | --- | --- | --- |
|  |  |  |  |  | Glide | Gold | rDock | Best Z-score | Mean Z-score |
| SMN2 Pre-mRNA | Groove Binding | 1 | 6HMO | 25 | 19.79 | 25.21 | 26.12 | 14.16 | 10.94 |
| Tau mRNA stemloop |  | 5 | 1EI2 | 7 | 14.96 | 16.21 | **31.43** | 17.02 | <u>20.53</u> |
| CAG Repeat |  | 7 | 7D12 | 11 | 22.98 | 25.05 | 18.69 | **32.94** | 16.04 |
| 16s rRNA A-Site (1)* |  | 50 | 2PWT | 66 | 21.53 | 22.60 | 26.14 | 19.63 | 17.99 |
| 16s rRNA A-Site (2)* |  | 12 | 5ZEI | 6 | 33.34 | **40.57** | 33.52 | 28.90 | <u>38.83</u> |
| HIV-1 FSS |  | 5 | 2JUK | 9 | **27.85** | 23.50 | 23.75 | <u>26.15</u> | 12.68 |
| HIV-TAR |  | 12 | 1AKX | 84 | 19.88 | **24.70** | 19.59 | 14.82 | 6.83 |
| IFA RNA promoter |  | 1 | 2LWK | 8 | 18.67 | 5.72 | **25.51** | 9.21 | <u>26.45</u> |
| Theophylline Aptamer | Nucleotide-Like Binding | 7 | 8D28 | 5 | 37.04 | 25.61 | 39.53 | 32.77 | 38.99 |
| Adenine Riboswitch |  | 16 | 4TZX | 17 | 53.48 | 33.07 | **60.73** | 48.49 | <u>57.65</u> |
| Guanine Riboswitch |  | 12 | 4FE5 | 33 | 45.14 | 30.07 | 48.81 | 40.06 | 46.63 |
| NAD<sup>+</sup> Riboswitch |  | 20 | 6TF0 | 9 | 30.92 | **48.91** | 36.12 | 31.85 | 36.06 |
| PreQ1 Riboswitch |  | 46 | 6E1W | 12 | 55.03 | 37.82 | 56.18 | <u>58.77</u> | 57.83 |
| SAM-I Riboswitch |  | 14 | 5FJC | 6 | 55.94 | 60.70 | 29.79 | 52.34 | 59.93 |
| SAM/SAH Riboswitch |  | 25 | 6LY5 | 6 | 27.37 | **45.92** | 24.33 | <u>39.48</u> | <u>43.78</u> |
| THF-I Riboswitch |  | 7 | 4LVW | 16 | 32.68 | 13.84 | 30.86 | 23.20 | <u>33.96</u> |
| THF-II Riboswitch |  | 3 | 7WIF | 11 | **33.34** | 16.19 | 23.4 | <u>27.12</u> | <u>31.92</u> |
| TPP Riboswitch |  | 21 | 2GDI | 12 | 47.65 | 31.17 | 45.87 | <u>47.25</u> | <u>51.34</u> |
| ZTP Riboswitch |  | 4 | 5BTP | 7 | 38.33 | 19.42 | **48.24** | <u>49.74</u> | <u>50.80</u> |
| Corn Aptamer | Intercalating | 5 | 5BJO | 6 | **46.05** | 33.12 | 32.10 | <u>36.81</u> | <u>39.67</u> |
| Pepper Aptamer |  | 12 | 7EOG | 8 | 63.88 | 60.38 | 57.74 | <u>62.73</u> | <u>66.51</u> |
| FMN Riboswitch |  | 33 | 5C45 | 25 | 38.09 | 38.76 | 24.83 | 34.87 | **46.13** |
| EV71 IRES |  | 8 | 6XB7 | 5 | 18.65 | 20.47 | **60.88** | <u>46.21</u> | <u>28.07</u> |
| HCV IRES |  | 1 | 2KTZ | 20 | **34.47** | 27.49 | 19.02 | 23.26 | <u>38.72</u> |
| Lysine Riboswitch | Misc. | 18 | 3DIL | 6 | 8.21 | **61.15** | 49.36 | <u>59.89</u> | <u>55.53</u> |
| Best in Targets# |  |  |  |  | 4 | 5 | 5 | 1 | 1 |
| Above the Median of Three Programs |  |  |  |  | - |  |  | 10 | 13 |

For the three methods, the best-performing one (with logAUC at least 10% better than the median) for each RNA target was shown in bold.
For consensus enrichment, the method with results 10% better than the best or median out of the three individual programs was shown in bold and underlined.
\*(1) and (2) stand for typical eukaryotic and prokaryotic sequences of 16s rRNA A-site.

Comparable performance between both the smaller and larger grid sizes was observed for 18, 11, and 11 RNA targets using Glide, Gold, and rDock, respectively. For instance, with both the grid sizes, the ligand enrichment from Glide, Gold and rDock for the pepper aptamer are similar, with logAUC of 63.88, 60.38, 57.74 and 65.38, 62.33, 53.09 for the smaller and larger grid size, respectively. Overall, Glide’s performance remained consistent in both grid sizes, while Gold and rDock shown varied results. Notably, rDock performed better with the smaller grid, underscoring the importance of choosing appropriate grid parameters in RNA-ligand docking. We note that the smaller grid improves ligand enrichment and pose prediction accuracy, particularly when using rDock.

To evaluate the impact of grid size on ligand enrichment, a paired t-test was also conducted, revealing a marginally significant overall difference between the two settings (p = 0.051, two-tailed). Among the three docking methods, rDock showed a statistically significant improvement with the smaller grid (p = 0.028). In contrast, Glide and Gold showed no significant differences (p = 0.553 and 0.234, respectively). As the smaller grid produced results that were either comparable to or better than those from the larger grid, we chose to prioritize the results generated with the small grid for subsequent analyses.

### Comparing Ligand Enrichment across Docking Programs and RNA Targets

We evaluated the ligand enrichment performance of Glide, Gold, and rDock programs to identify the best performing programs for docking against RNA targets. As highlighted in Table 2, Glide, Gold, and rDock outperformed the other two programs in four, five, and five RNA targets, respectively. For example, Glide performed best on the corn aptamer and HCV IRES, with logAUC values of 46.05, and 34.47, respectively. Gold showed superior enrichment for the lysine and NAD+ riboswitches, with logAUC values of 61.15 and 48.91, respectively. rDock performed best for the ZTP riboswitch and EV71 IRES, with logAUC values of 48.24 and 60.88, respectively. For the remaining eleven RNA targets, all three programs exhibited comparable performance. In some cases, all three methods demonstrated strong ligand enrichment, as observed for the pepper aptamer, with logAUC values of 63.88, 60.38, and 57.74 for Glide, Gold, and rDock, respectively. Conversely, all three programs performed poorly for certain targets, such as the eukaryotic 16s rRNA A-site, with logAUC values of 22.98, 25.05, and 18.69 for Glide, Gold, and rDock, respectively. In other instances, two programs outperformed the third, as seen for the guanine and SAM-I riboswitches with logAUC of 45.14, 30.07, 48.81 and 55.94, 60.70, 29.79 using Glide, Gold and rDock, respectively. Overall, the three programs did not show statistically significant differences in performance (p = 0.54, two-way ANOVA); however, their effectiveness varied depending on the RNA targets (p < 0.05, two-way ANOVA).

While the overall performance between three methods were not significant, the performance varied significantly for RNA targets associated with different modes of binding. As shown in Table 2, all three methods achieved better ligand enrichment for RNA targets associated with intercalating or nucleotide-like ligand binding, whereas groove binding RNA targets posed the greatest challenge for RNA-ligand docking. For the five RNA targets with intercalating binding, 13 of 15 docking runs exceeded the logAUC threshold of 24.5 that is twice the enrichment of random selection. Glide achieved the best ligand enrichment for the corn aptamer and HCV IRES, with logAUC values of 46.05 and 34.47, respectively. In contrast, rDock significantly outperformed the other methods for EV-71 IRES, achieving a logAUC of 60.88. For the 11 RNA targets associated with nucleotide-like binding mode, performance was notably higher, with 28 out of 33 docking runs exceeding two-fold enrichment of random selection. As highlighted in Table 2, Gold demonstrated superior performance against NAD+ and SAM-SAH riboswitches (logAUC of 48.91 and 45.92), while rDock does best for adenine and ZTP riboswitches (logAUC of 60.73 and 48.24).

In contrast, docking performance was substantially worse for the eight RNA targets associated with groove binding, with only 13 of 24 docking runs yielding two-fold enrichment of random selection. Among these, rDock performed best for tau pre-mRNA and the influenza RNA promoter (logAUC of 31.43 and 25.51), while Gold was more effective for the prokaryotic 16S rRNA A-site and HIV TAR (logAUC of 40.57 and 24.70). Glide only showed best performance against one groove binding RNA target HIV FSS (logAUC of 27.85). Beyond the three categories based on ligand binding modes, lysine riboswitch exhibited better enrichment with Gold and rDock than Glide (logAUC of 8.21, 61.15, and 49.36, for Glide, Gold, and rDock, respectively).

These findings highlight the importance of using the most appropriate docking method for specific RNA target properties to improve predictive accuracy in RNA-ligand docking studies. While all three methods performed well for RNA targets with intercalating and nucleotide-like binding modes, their effectiveness varied for individual targets. RNA targets associated with groove binding typically lack a well-defined binding pocket, which may result in lower docking performance. For example, the ligands for RNA targets with intercalating and nucleotide-like binding modes have ligands that mostly intercalate between base pairs and form hydrogen bonds within a structured cavity. In contrast, ligands for groove binding RNA targets interact along the RNA helix with hydrogen bonds and electrostatic interactions, without deeply embedding within the structure, which could lead to more ambiguous modes of binding. This suggests that current docking algorithms may struggle to model interactions for groove binding ligands accurately due to limitations in scoring functions and the treatment of RNA flexibility. To improve the design of groove binding ligands, future studies could develop more refined scoring functions that better capture hydrogen bonding and electrostatic interactions, along with more comprehensive sampling of RNA conformations.

### Consensus Ligand Enrichment across Docking Programs

To evaluate whether combining results from multiple docking programs enhances performance, we calculated consensus enrichment using both the best and mean Z-score for each ligand and decoy. Both approaches showed some improvement over individual docking methods. Consensus enrichment based on the best Z-score and the mean Z-score exceeded the median logAUC of the three programs by at least 10% in 10 and 13 out of 25 RNA targets, respectively. However, when compared to the best individual logAUC, both approaches achieved a 10% improvement in only one case: the best Z-score approach for the CAG repeat (logAUC of 32.94) and the mean Z-score approach for the FMN riboswitch (logAUC of 46.13). Notably, for 6 and 14 targets, the difference between the consensus enrichment obtained from the best Z-scores or mean Z-scores and the enrichment of the best docking method is less than 10% of the latter, respectively. Therefore, the mean Z-score method yields slightly but consistently better performance than the best Z-score method. Based on these results, integrating multiple docking methods can result in suboptimal ligand enrichment that is consistently better when compared to using them alone. This is useful in practical situations when the most effective technique is uncertain.

Averaging the scores can make the performance more robust; by taking the mean, outlier scores from any single method are balanced by the others, producing predictions that are more consistent across targets. This effect is evident for adenine riboswitch, where the ligand enrichment for three methods varied significantly, with logAUC of 53.48, 33.07, and 60.73, respectively from Glide, Gold, and rDock. Despite Gold’s relatively poor performance, the consensus enrichment based on the mean Z-score reached 57.65, which is not only within 10% of the best-performing method (rDock) but also significantly outperformed the result derived from the best Z-score alone (48.49).In contrast, selecting the best individual score may occasionally capture an exceptional result, but it is more susceptible to variability and may not generalize well across different RNA targets. This suggests that consensus scoring based on the mean Z-score provides a reliable strategy for RNA-targeted virtual screening.

### Consensus over Multiple RNA Structures

In this study, we also evaluate whether combining results from multiple structures of each target enhances performance, by including two additional structures for six targets: X-ray structures for eukaryotic 16s rRNA A-Site, preQ1 riboswitch and TPP riboswitch, and NMR structures for tau pre-mRNA hairpin, EV71 IRES and HIV FSS. As described in the methods section, the selection of additional structures was based on RMSD from structural alignment of RNA backbone to maximize the RMSD values between the structures (Table S5). Consensus enrichment was calculated using the mean or the best docking score from each docking method across the three structures for each target (Table S6). Among 18 target-method combinations (6 targets x 3 docking methods), the consensus enrichment based on the mean score and the best score outperformed the median logAUC in 3 and 4 cases, respectively, but never exceeded the best individual logAUC by 10%. However, in 10 and 11 cases, the difference between the consensus enrichment derived from the best and mean scores, and the best individual logAUC is less than 10% of the latter, respectively. These results suggest that consensus enrichment across multiple structures can also yield suboptimal but stable ligand enrichment compared to that from an individual structure, alike consensus enrichment across multiple docking methods.

### Performance for different Chemotypes of Ligands

To investigate the distribution of molecular scaffolds in the dataset and determine whether ligand chemotypes affect docking performance, we conducted chemical clustering for RNA binders. A total of 432 RNA-binding molecules were clustered using the HDBSCAN algorithm (minimum cluster size of 20, minimum sample of 4, and clustering selection method is leaf), based on the uniform manifold approximation and projection (UMAP) embedding of the Morgan fingerprint Tanimoto distance matrix. The resulting ten clusters, comprising 406 ligands (26 outliers), were visualized using a scatterplot, which demonstrated well-defined groupings (Figure 3A). The overall scaffold distribution and representative chemotypes for each cluster are shown in Figure 3C and Table S7.

Notably, each cluster contained ligands from multiple RNA targets, indicating that structurally diverse scaffolds are not limited to specific RNA targets but can interact with a variety of RNA targets. Among the ten clusters, clusters 1, 2, and 3, which contain 54, 29, and 82 ligands, respectively, accounted for ∼ 40% of the dataset. As shown in Figure 3C, these three clusters are composed of well-known RNA-binding scaffolds that include aminoglycosides, nucleoside derivatives, and nucleotide derivatives whereas clusters 4– 10, each containing 27–47 ligands, possessed more structurally diverse scaffolds. Most molecules under clusters 4-10 contain aromatic carbocycles and were more structurally similar to each other than clusters 1–3. Chemotype distribution suggests that both well-characterized and diverse types of RNA binders are present among the clusters, indicating that while certain chemotypes are recurrent in RNA-ligand interactions, a broader range of chemical scaffolds also contribute to RNA recognition.

We analyzed the percentile rank distributions of RNA binders from three docking methods: Glide, Gold, and rDock in order to further investigate variations in ligand enrichment among scaffolds (Figure 3B). RNA binders in clusters 1, 2, 3, and 6 were ranked significantly higher, whereas those in other clusters showed a lower or more diverse distribution of ranks. Statistical analysis identified docking method-specific trends in clusters 1, 3, 4, 5, 7, 10 (p < 0.05, Kruskal-Walis test). For RNA binders in clusters 1 and 7, rDock worked best, while Gold was more successful in clusters 5 and 10 but less successful in cluster 3. Furthermore, Gold performed considerably better than Glide and somewhat better than rDock in cluster 4, indicating that some ligand chemotypes might be less suited for particular docking method(s). These findings demonstrate how ligand chemotype composition affects docking performance and emphasizes the significance of docking method choice against the target RNA with respect to the characteristics of known RNA binders.

## CONCLUSION

In this study, we compiled a dataset of 25 RNA targets with experimental structures and corresponding ligands to systematically assess the performance of three docking programs for RNA-ligand docking screens. Adjustment of the grid size parameter revealed that a smaller grid size enhanced native ligand binding geometry prediction accuracy across all methods, while rDock also showed improved ligand enrichment during virtual screening under this setting. This is because the smaller grid gives a more constrained binding cavity that improves the accuracy of pose sampling in docking programs. These observations highlight the importance of accurately defined binding sites in future RNA-ligand docking studies.

While performance differences among the docking programs were not significant enough for all the 25 RNA targets, the performances across the different classes of RNA targets were substantially different. We observed all three docking programs struggled against groove binding RNA targets. In contrast, their performance was notably better for those associated with intercalating and nucleic acid-like binding, where the binding pockets are more defined and structured, allowing for better sampling and scoring.

Consensus enrichment analysis based on different docking methods and receptor conformations yielded stable and slightly superior performance compared to individual docking methods. Specifically, consensus enrichment across multiple docking methods outperformed the median method for 13 of the 25 RNA targets, while consensus enrichment across multiple structures produced performance comparable to the best individual structure for 11 of the 18 target-method combinations. Between the two approaches for calculating consensus—mean score and best score—the mean approach showed slightly better overall consistency, suggesting that averaging scores may help reduce the variability across docking methods. In contrast, the best-score approach may occasionally capture favorable binding poses but is more sensitive to outliers. These observations indicate that while consensus strategies do not substantially improve enrichment, they may offer practical benefits in stabilizing performance across diverse RNA targets.

Clustering of the ligands by chemotype further revealed method-specific and chemotype specific biases. Among ten ligand clusters, six showed distinct performance trends across the three docking programs. For instance, rDock ranked ligands in clusters 1 and 7 significantly higher, suggesting better compatibility with aminoglycoside-like and certain aromatic scaffolds. Gold performed well in clusters 5 and 10, which included diverse aromatic carbocycles, but was notably less effective in cluster 3, enriched with nucleotide-like ligands. In contrast, Glide consistently underperformed in several clusters, particularly in clusters 4 and 10. Glide This suggest that the choice of docking method may favor certain ligand chemotypes or scaffolds.

Overall, our results indicate that current docking programs can be applied to RNA targets with known experimental structures, though their effectiveness will depend on the target class and ligand properties. To better capture the dynamic nature of RNA and enhance docking-based virtual screening performance for RNA-targeted drug discovery, future studies could focus on refining RNA-ligand scoring functions (e.g. optimizing electrostatic treatments to better capture groove binding interactions) and incorporating RNA flexibility through ensemble docking, RNA structure modeling, and molecular dynamics simulations.

## ACKNOWLEDGEMENTS

The author would like to thank Dr. Ravi Kumar Verma and Dr. Mandar Kulkarni for their support and valuable feedback.

## AUTHOR CONTRIBUTIONS

Conceptualization: H.F., R.G.H., and C.L.L.C.; Data collection: L.D. and W.K.; Computation: L.D. and W.K.; Writing and discussion: W.K., L.D., C.K.J., H.F., R.G.H., and C.L.L.C.; Funding acquisition: H.F., R.G.H., and C.L.L.C.

## SUPPLEMENTARY DATA

Supplementary data are available at NAR online.

## DATA AVAILABILITY

All data are available in the main text or supplementary data.

