## Supplemental Table (S4-S7) and Supplemental Figures for "Evaluating Molecular Docking Programs for RNA-Targeted Ligand Screening: Influence of Binding Modes and Ligand Types"

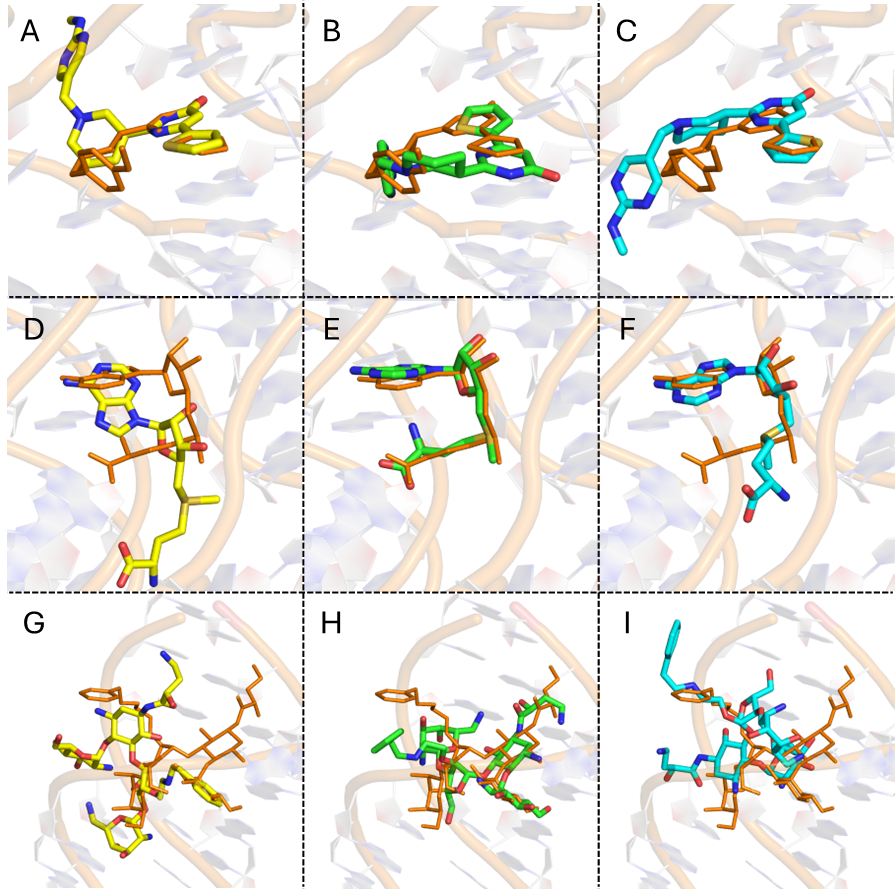

**Figure S1**. The experimental structures (orange) and binding poses predicted by Glide (yellow), Gold (green) and rDock (cyan) of the native ligands from three classes of RNA targets with different binding modes. (A, B, C) For RNA targets associated with intercalating binding: FMN riboswitch (PDB: 5C45). (D, E, F) For RNA targets associated with nucleotide-like binding: SAM-I riboswitch (PDB: 5FJC). (G, H, I) For RNA targets associated with groove binding: 16s rRNA A-site (PDB: 2PWT).

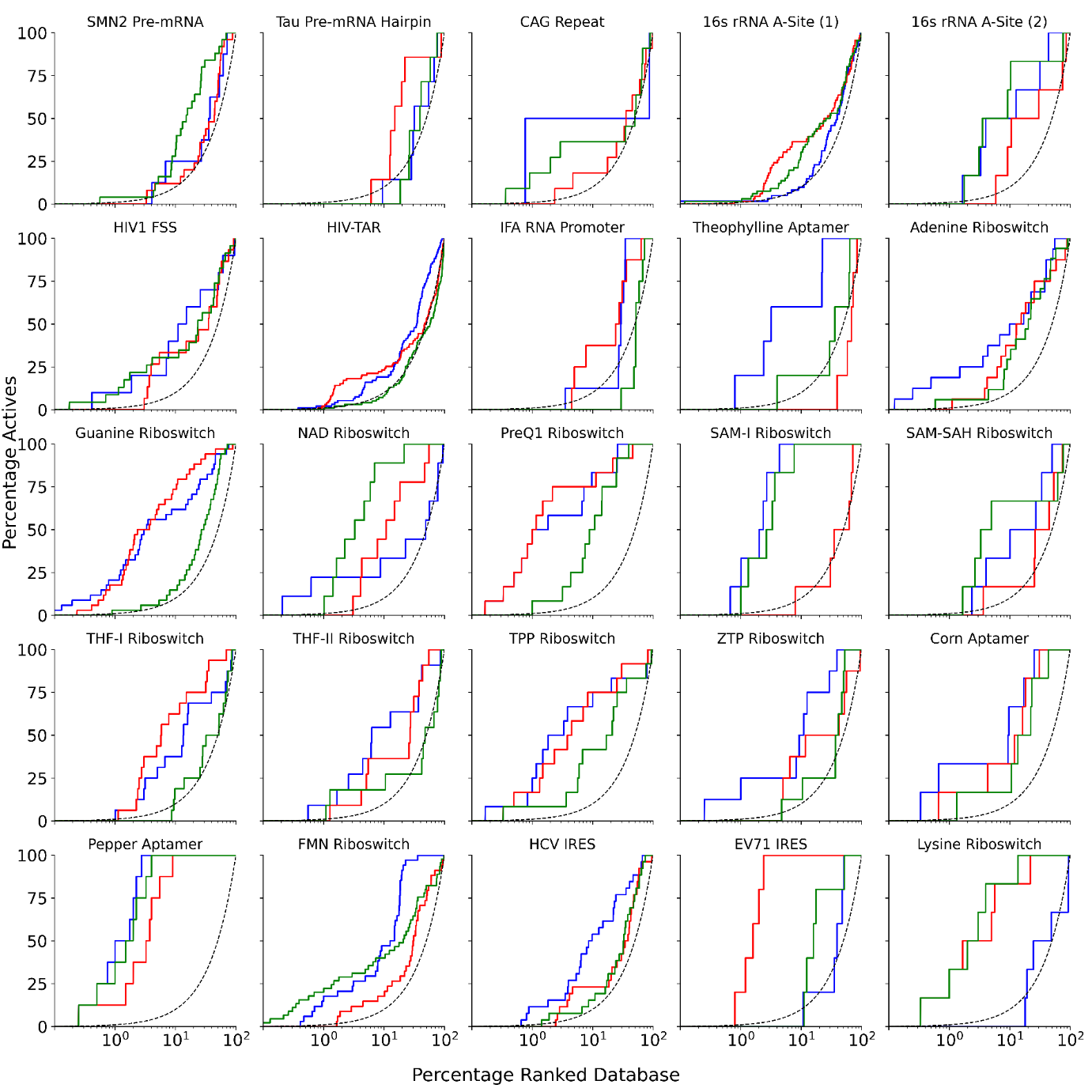

**Figure S2**. Enrichment curves for Glide (blue), Gold (green) and rDock (red), and random selection (dotted line) of the 25 RNA targets with larger grid size for docking.

| **Table S4**. Ligand enrichment results (logAUC) for Glide, Gold, and rDock against 25 representative PDB structures, with smaller and larger grid sizes applied for docking. | | | | | | | | |
| --- | --- | --- | --- | --- | --- | --- | --- | --- |
| Target | Category | Representative PDB | Larger Grid Size | | | Smaller Grid Size | | |
|  |  |  | Glide | Gold | rDock | Glide | Gold | rDock |
| Artificial Aptamers | | | | | | | | |
| Corn Aptamer | Intercalating | 5BJO | 44.75 | 29.79 | 36.12 | 46.05 | 33.12 | 32.10 |
| Pepper Aptamer | Intercalating | 7EOG | 65.38 | 62.33 | 53.09 | 63.88 | 60.38 | 57.74 |
| Theophylline Aptamer | Nucleotide-Like | 8D28 | 43.36 | 18.50 | 31.81 | 37.04 | 25.61 | 39.53 |
| Human RNA | | | | | | | | |
| SMN2 Pre-mRNA | Groove Binding | **6HMO** | 19.32 | 28.12 | 18.11 | 19.97 | 25.21 | 26.12 |
| Tau Pre-mRNA | Groove Binding | **1EI2** | 15.02 | 14.53 | 25.00 | 14.96 | 16.21 | 31.43 |
| CAG Repeat | Groove Binding | **7D12** | 34.12 | 28.37 | 18.37 | 22.98 | 25.05 | 18.69 |
| Bacterial Ribosomal RNA | | | | | | | | |
| 16s rRNA A-Site (1)* | Groove Binding | 2PWT | 20.86 | 23.53 | 25.93 | 21.53 | 22.60 | 26.14 |
| 16s rRNA A-Site (2)* | Groove Binding | 5ZEI | 35.63 | 38.05 | 21.95 | 33.34 | 40.57 | 33.52 |
| Bacterial Riboswitch | | | | | | | | |
| Adenine Riboswitch | Nucleotide-Like | 4TZX | 38.47 | 26.07 | 27.87 | 53.22 | 33.49 | 39.40 |
| Guanine Riboswitch | Nucleotide-Like | 4FE5 | 45.37 | 22.59 | 47.95 | 45.05 | 30.07 | 48.81 |
| Lysine Riboswitch | Misc | 3DIL | 13.54 | 55.34 | 52.56 | 8.21 | 61.15 | 49.36 |
| NAD+ Riboswitch | Nucleotide-Like | 6TF0 | 27.43 | 48.68 | 31.46 | 30.92 | 48.91 | 36.12 |
| PreQ1 Riboswitch | Nucleotide-Like | 6E1W | 55.73 | 34.66 | 58.84 | 56.32 | 36.74 | 57.68 |
| SAM-I Riboswitch | Nucleotide-Like | 5FJC | 57.88 | 52.65 | 14.40 | 55.94 | 60.70 | 29.79 |
| SAM-SAH Riboswitch | Nucleotide-Like | 6LY5 | 29.72 | 35.88 | 18.67 | 27.37 | 45.92 | 24.33 |
| THF-I Riboswitch | Nucleotide-Like | 4LVW | 29.65 | 15.66 | 37.80 | 32.68 | 13.84 | 30.86 |
| THF-II Riboswitch | Nucleotide-Like | 7WIF | 35.14 | 19.37 | 27.58 | 33.34 | 16.19 | 23.40 |
| TPP Riboswitch | Nucleotide-Like | 2GDI | 47.53 | 30.09 | 45.32 | 47.65 | 31.17 | 45.87 |
| ZTP Riboswitch | Nucleotide-Like | 5BTP | 39.68 | 18.57 | 23.63 | 38.33 | 19.42 | 48.24 |
| FMN Riboswitch | Intercalating | 5C45 | 38.31 | 37.94 | 21.48 | 38.09 | 38.76 | 24.83 |
| Viral RNA | | | | | | | | |
| EV71 IRES | Intercalating | **6XB7** | 16.29 | 24.61 | 60.88 | 18.65 | 20.47 | 60.88 |
| HCV IRES | Intercalating | **2KTZ** | 33.89 | 28.49 | 21.59 | 34.47 | 27.49 | 19.02 |
| HIV FSS | Groove Binding | **2JUK** | 30.93 | 31.34 | 23.83 | 27.85 | 23.50 | 23.75 |
| HIV-TAR | Groove Binding | **1AKX** | 21.63 | 24.96 | 20.89 | 20.87 | 24.70 | 19.59 |
| IFA RNA promoter | Groove Binding | **2LWK** | 21.09 | 9.36 | 25.36 | 18.67 | 5.72 | 25.51 |
| The RNA targets with NMR structure as representative structure are highlighted.  *: (1) and (2) stand for typical eukaryotic and prokaryotic sequences of 16s rRNA A-site. | | | | | | | | |

**Table S5**. RMSD values from superposition of cavity residues (within 6 Å of the native ligand) for selected X-Ray structures and NMR conformers.

| **X-Ray Structures: 16s rRNA (eukaryotic)** | | | | |
| --- | --- | --- | --- | --- |
| Cluster Index | PDB | 1LC4 | 1YRJ | 4F8U |
| 81 | 1LC4 | 0.000 | 1.520 | 2.466 |
| 81 | 1YRJ | 1.52 | 0.000 | 2.857 |
| 81 | 2ESI | 0.668 | 1.813 | 1.953 |
| 81 | 2O3X | 0.928 | 1.661 | 2.629 |
| 81 | 4F8U | 2.466 | 2.857 | 0.000 |
| 81 | 4P20 | 0.34 | 1.679 | 2.482 |
| 123 | 1J7T | 0.700 | 1.534 | 2.609 |
| 123 | 1MWL | 0.747 | 1.402 | 2.738 |
| 123 | 2BE0 | 1.206 | 1.590 | 3.747 |
| 123 | 2BEE | 1.171 | 1.605 | 3.678 |
| 123 | 2ESJ | 0.900 | 1.348 | 2.907 |
| 123 | 2ET3 | 1.038 | 1.283 | 3.074 |
| 123 | 2ET4 | 0.628 | 1.392 | 2.923 |
| 123 | 2ET5 | 0.732 | 1.563 | 2.912 |
| 123 | 2ET8 | 1.040 | 1.761 | 3.000 |
| 123 | 2F4S | 1.201 | 1.799 | 2.743 |
| 123 | 2F4U | 1.125 | 1.820 | 3.006 |
| 123 | 2PWT | 1.277 | 2.531 | 3.444 |
| 123 | 3S4P | 0.994 | 1.711 | 2.805 |

| **X-Ray Structures: PreQ1 Riboswitch** | | | | | |
| --- | --- | --- | --- | --- | --- |
| Cluster Index | PDB | 6E1S | 6E1U | 6E1W | 7E9E |
| 110 | 6E1S | 0.000 | 0.311 | 1.304 | 1.256 |
| 110 | 6E1U | 0.311 | 0.000 | 1.176 | 1.088 |
| 110 | 6E1W | 1.304 | 1.176 | 0.000 | 1.008 |
| 110 | 7E9E | 1.256 | 1.088 | 1.008 | 0.000 |

| **X-Ray Structures: TPP Riboswitch** | | | | | | | | | | | | | | | | | | | | | |
| --- | --- | --- | --- | --- | --- | --- | --- | --- | --- | --- | --- | --- | --- | --- | --- | --- | --- | --- | --- | --- | --- |
| Cluster Index | PDB | 2GDI | 2HOJ | 2HOK | 2HOL | 2HOM | 2HOO | 2HOP | 4NYA | 4NYB | 4NYC | 4NYD | 4NYG | 7TD7 | 7TDA | 7TDB | 7TDC | 7TZR | 7TZS | 7TZT | 7TZU |
| 34 | 2GDI | 0.000 | 0.410 | 1.151 | 0.425 | 0.574 | 1.089 | 1.247 | 0.396 | 1.030 | 0.852 | 0.593 | 1.241 | 0.720 | 0.533 | 0.744 | 0.766 | 1.010 | 0.873 | 0.903 | 0.959 |
| 34 | 2HOJ | 0.410 | 0.000 | 1.141 | 0.174 | 0.422 | 0.956 | 1.082 | 0.486 | 0.854 | 0.628 | 0.381 | 1.182 | 0.640 | 0.380 | 0.511 | 0.544 | 0.933 | 0.839 | 0.776 | 0.831 |
| 34 | 2HOK | 1.151 | 1.141 | 0.000 | 0.391 | 0.355 | 0.768 | 1.001 | 0.598 | 0.494 | 0.437 | 0.433 | 1.140 | 0.571 | 0.399 | 0.317 | 0.348 | 0.690 | 0.808 | 0.730 | 0.765 |
| 34 | 2HOL | 0.425 | 0.174 | 0.391 | 0.000 | 0.408 | 0.949 | 1.087 | 0.502 | 0.937 | 0.723 | 0.396 | 1.182 | 0.664 | 0.482 | 0.595 | 0.622 | 1.014 | 0.900 | 0.812 | 0.827 |
| 34 | 2HOM | 0.574 | 0.422 | 0.355 | 0.408 | 0.000 | 0.701 | 1.041 | 0.560 | 0.811 | 0.638 | 0.413 | 1.058 | 0.601 | 0.530 | 0.500 | 0.545 | 0.922 | 0.809 | 0.710 | 0.773 |
| 34 | 2HOO | 1.089 | 0.956 | 0.768 | 0.949 | 0.701 | 0.000 | 0.820 | 0.902 | 0.932 | 0.866 | 0.742 | 0.974 | 0.650 | 0.922 | 0.886 | 0.927 | 1.085 | 1.041 | 0.626 | 0.615 |
| 34 | 2HOP | 1.247 | 1.082 | 1.001 | 1.087 | 1.041 | 0.820 | 0.000 | 5.529 | 5.428 | 5.563 | 5.449 | 5.277 | 5.452 | 5.506 | 5.529 | 5.568 | 5.526 | 5.474 | 5.319 | 5.184 |
| 34 | 4NYA | 0.396 | 0.486 | 0.598 | 0.502 | 0.560 | 0.902 | 5.529 | 0.000 | 0.885 | 0.720 | 0.522 | 1.132 | 0.681 | 0.487 | 0.627 | 0.679 | 0.886 | 0.706 | 0.828 | 0.904 |
| 34 | 4NYB | 1.030 | 0.854 | 0.494 | 0.937 | 0.811 | 0.932 | 5.428 | 0.885 | 0.000 | 0.565 | 0.295 | 1.050 | 0.589 | 0.724 | 0.608 | 0.633 | 0.489 | 0.580 | 0.857 | 0.927 |
| 34 | 4NYC | 0.852 | 0.628 | 0.437 | 0.723 | 0.638 | 0.866 | 5.563 | 0.720 | 0.565 | 0.000 | 0.341 | 1.168 | 0.562 | 0.458 | 0.410 | 0.406 | 0.699 | 0.782 | 0.676 | 0.768 |
| 34 | 4NYD | 0.593 | 0.381 | 0.433 | 0.396 | 0.413 | 0.742 | 5.449 | 0.522 | 0.295 | 0.341 | 0.000 | 1.123 | 0.613 | 0.418 | 0.493 | 0.534 | 0.902 | 0.798 | 0.684 | 0.721 |
| 34 | 4NYG | 1.241 | 1.182 | 1.140 | 1.182 | 1.058 | 0.974 | 5.277 | 1.132 | 1.050 | 1.168 | 1.123 | 0.000 | 1.077 | 1.233 | 1.237 | 1.266 | 1.224 | 1.169 | 0.968 | 0.969 |
| 34 | 7TD7 | 0.720 | 0.640 | 0.571 | 0.664 | 0.601 | 0.650 | 5.452 | 0.681 | 0.589 | 0.562 | 0.613 | 1.077 | 0.000 | 4.363 | 4.395 | 4.359 | 0.694 | 0.912 | 0.538 | 0.581 |
| 34 | 7TDA | 0.533 | 0.380 | 0.399 | 0.482 | 0.530 | 0.922 | 5.506 | 0.487 | 0.724 | 0.458 | 0.418 | 1.233 | 4.363 | 0.000 | 0.309 | 0.358 | 0.779 | 0.745 | 0.730 | 0.802 |
| 34 | 7TDB | 0.744 | 0.511 | 0.317 | 0.595 | 0.500 | 0.886 | 5.529 | 0.627 | 0.608 | 0.410 | 0.493 | 1.237 | 4.395 | 0.309 | 0.000 | 0.186 | 0.651 | 0.671 | 0.737 | 0.811 |
| 34 | 7TDC | 0.766 | 0.544 | 0.348 | 0.622 | 0.545 | 0.927 | 5.568 | 0.679 | 0.633 | 0.406 | 0.534 | 1.266 | 4.359 | 0.358 | 0.186 | 0.000 | 0.691 | 0.724 | 0.803 | 0.868 |
| 34 | 7TZR | 1.010 | 0.933 | 0.690 | 1.014 | 0.922 | 1.085 | 5.526 | 0.886 | 0.489 | 0.699 | 0.902 | 1.224 | 0.694 | 0.779 | 0.651 | 0.691 | 0.000 | 0.435 | 0.892 | 1.025 |
| 34 | 7TZS | 0.873 | 0.839 | 0.808 | 0.900 | 0.809 | 1.041 | 5.474 | 0.706 | 0.580 | 0.782 | 0.798 | 1.169 | 0.912 | 0.745 | 0.671 | 0.724 | 0.435 | 0.000 | 0.909 | 1.043 |
| 34 | 7TZT | 0.903 | 0.776 | 0.730 | 0.812 | 0.710 | 0.626 | 5.319 | 0.828 | 0.857 | 0.676 | 0.684 | 0.968 | 0.538 | 0.730 | 0.737 | 0.803 | 0.892 | 0.909 | 0.000 | 0.377 |
| 34 | 7TZU | 0.959 | 0.831 | 0.765 | 0.827 | 0.773 | 0.615 | 5.184 | 0.904 | 0.927 | 0.768 | 0.721 | 0.969 | 0.581 | 0.802 | 0.811 | 0.868 | 1.025 | 1.043 | 0.377 | 0.000 |

| **NMR Conformers: Tau Pre-mRNA Hairpin** | | | | | | | | | | | | | | | | | |
| --- | --- | --- | --- | --- | --- | --- | --- | --- | --- | --- | --- | --- | --- | --- | --- | --- | --- |
| 1EI2 | C1 | C2 | C3 | C4 | C5 | C6 | C7 | C8 | C9 | C10 | C11 | C12 | C13 | C14 | C15 | C16 | C17 |
| C1 | 0.000 | 1.316 | 3.479 | 4.883 | 3.943 | 4.076 | 3.147 | 4.407 | 4.655 | 3.709 | 4.894 | 5.944 | 4.505 | 3.268 | 6.949 | 4.266 | 5.441 |
| C2 | 1.316 | 0.000 | 3.229 | 4.199 | 3.527 | 3.509 | 2.843 | 3.918 | 3.956 | 3.070 | 4.309 | 5.182 | 3.925 | 3.273 | 6.245 | 3.883 | 4.857 |
| C3 | 3.479 | 3.229 | 0.000 | 3.287 | 2.915 | 2.322 | 2.464 | 3.144 | 3.439 | 3.509 | 1.702 | 4.765 | 2.780 | 2.961 | 6.927 | 3.489 | 4.589 |
| C4 | 4.883 | 4.199 | 3.287 | 0.000 | 1.476 | 2.273 | 1.913 | 1.985 | 1.796 | 2.977 | 3.286 | 4.476 | 3.941 | 2.715 | 6.234 | 2.291 | 2.867 |
| C5 | 3.943 | 3.527 | 2.915 | 1.476 | 0.000 | 2.754 | 1.600 | 2.003 | 1.995 | 2.898 | 3.099 | 4.395 | 3.523 | 2.101 | 6.461 | 2.074 | 2.986 |
| C6 | 4.076 | 3.509 | 2.322 | 2.273 | 2.754 | 0.000 | 2.095 | 1.874 | 2.333 | 2.834 | 2.169 | 3.355 | 2.522 | 2.207 | 5.771 | 3.272 | 3.273 |
| C7 | 3.147 | 2.843 | 2.464 | 1.913 | 1.600 | 2.095 | 0.000 | 1.193 | 1.789 | 2.296 | 2.478 | 2.486 | 2.729 | 1.978 | 4.794 | 1.615 | 1.937 |
| C8 | 4.407 | 3.918 | 3.144 | 1.985 | 2.003 | 1.874 | 1.193 | 0.000 | 2.011 | 2.221 | 2.895 | 3.210 | 3.185 | 1.746 | 5.762 | 1.591 | 2.321 |
| C9 | 4.655 | 3.956 | 3.439 | 1.796 | 1.995 | 2.333 | 1.789 | 2.011 | 0.000 | 2.385 | 3.372 | 3.376 | 3.128 | 1.796 | 5.790 | 2.780 | 2.610 |
| C10 | 3.709 | 3.070 | 3.509 | 2.977 | 2.898 | 2.834 | 2.296 | 2.221 | 2.385 | 0.000 | 3.860 | 2.701 | 2.769 | 2.953 | 4.031 | 3.452 | 3.419 |
| C11 | 4.894 | 4.309 | 1.702 | 3.286 | 3.099 | 2.169 | 2.478 | 2.895 | 3.372 | 3.860 | 0.000 | 4.666 | 3.110 | 2.688 | 6.654 | 3.119 | 4.043 |
| C12 | 5.944 | 5.182 | 4.765 | 4.476 | 4.395 | 3.355 | 2.486 | 3.210 | 3.376 | 2.701 | 4.666 | 0.000 | 3.349 | 2.412 | 4.596 | 4.441 | 3.196 |
| C13 | 4.505 | 3.925 | 2.780 | 3.941 | 3.523 | 2.522 | 2.729 | 3.185 | 3.128 | 2.769 | 3.110 | 3.349 | 0.000 | 3.134 | 5.575 | 4.101 | 4.271 |
| C14 | 3.268 | 3.273 | 2.961 | 2.715 | 2.101 | 2.207 | 1.978 | 1.746 | 1.796 | 2.953 | 2.688 | 2.412 | 3.134 | 0.000 | 5.076 | 2.402 | 2.342 |
| C15 | 6.949 | 6.245 | 6.927 | 6.234 | 6.461 | 5.771 | 4.794 | 5.762 | 5.790 | 4.031 | 6.654 | 4.596 | 5.575 | 5.076 | 0.000 | 6.597 | 4.287 |
| C16 | 4.266 | 3.883 | 3.489 | 2.291 | 2.074 | 3.272 | 1.615 | 1.591 | 2.780 | 3.452 | 3.119 | 4.441 | 4.101 | 2.402 | 6.597 | 0.000 | 3.041 |
| C17 | 5.441 | 4.857 | 4.589 | 2.867 | 2.986 | 3.273 | 1.937 | 2.321 | 2.610 | 3.419 | 4.043 | 3.196 | 4.271 | 2.342 | 4.287 | 3.041 | 0.000 |

| **NMR Conformers: EV71 IRES** | | | | | | | | | | |
| --- | --- | --- | --- | --- | --- | --- | --- | --- | --- | --- |
| 6XB7 | C1 | C2 | C3 | C4 | C5 | C6 | C7 | C8 | C9 | C10 |
| C1 | 0.000 | 4.655 | 4.840 | 4.596 | 7.461 | 2.775 | 3.199 | 2.733 | 5.764 | 7.101 |
| C2 | 4.655 | 0.000 | 6.120 | 5.582 | 6.549 | 4.333 | 2.648 | 4.020 | 4.321 | 5.317 |
| C3 | 4.840 | 6.120 | 0.000 | 7.007 | 3.105 | 5.568 | 6.039 | 5.383 | 5.599 | 4.106 |
| C4 | 4.596 | 5.582 | 7.007 | 0.000 | 7.252 | 5.468 | 7.256 | 4.948 | 7.359 | 5.828 |
| C5 | 7.461 | 6.549 | 3.105 | 7.252 | 0.000 | 7.606 | 7.168 | 6.960 | 6.745 | 4.903 |
| C6 | 2.775 | 4.333 | 5.568 | 5.468 | 7.606 | 0.000 | 2.402 | 0.963 | 5.629 | 6.316 |
| C7 | 3.199 | 2.648 | 6.039 | 7.256 | 7.168 | 2.402 | 0.000 | 2.438 | 4.817 | 4.929 |
| C8 | 2.733 | 4.020 | 5.383 | 4.948 | 6.960 | 0.963 | 2.438 | 0.000 | 6.051 | 6.644 |
| C9 | 5.764 | 4.321 | 5.599 | 7.359 | 6.745 | 5.629 | 4.817 | 6.051 | 0.000 | 3.489 |
| C10 | 7.101 | 5.317 | 4.106 | 5.828 | 4.903 | 6.316 | 4.929 | 6.644 | 3.489 | 0.000 |

| **NMR Conformers: HIV FSS** | | | | | | | | | | | | | | | | | | | | |
| --- | --- | --- | --- | --- | --- | --- | --- | --- | --- | --- | --- | --- | --- | --- | --- | --- | --- | --- | --- | --- |
| 2JUK | C1 | C2 | C3 | C4 | C5 | C6 | C7 | C8 | C9 | C10 | C11 | C12 | C13 | C14 | C15 | C16 | C17 | C18 | C19 | C20 |
| C1 | 0.000 | 1.788 | 0.972 | 2.856 | 1.117 | 1.721 | 1.257 | 1.075 | 1.213 | 1.320 | 1.255 | 1.058 | 1.761 | 1.406 | 1.979 | 2.415 | 0.897 | 0.930 | 1.245 | 1.777 |
| C2 | 1.788 | 0.000 | 1.807 | 1.302 | 2.016 | 1.904 | 2.048 | 1.894 | 1.450 | 1.412 | 1.210 | 1.674 | 1.029 | 1.431 | 3.015 | 4.176 | 1.972 | 1.657 | 1.757 | 1.382 |
| C3 | 0.972 | 1.807 | 0.000 | 2.774 | 1.486 | 1.967 | 1.284 | 1.120 | 1.500 | 1.407 | 1.368 | 1.105 | 2.029 | 1.640 | 2.259 | 2.386 | 0.974 | 0.787 | 1.068 | 2.072 |
| C4 | 2.856 | 1.302 | 2.774 | 0.000 | 2.153 | 1.823 | 2.279 | 2.450 | 1.691 | 1.630 | 1.561 | 2.077 | 1.245 | 1.836 | 3.411 | 3.205 | 2.627 | 2.218 | 1.958 | 1.383 |
| C5 | 1.117 | 2.016 | 1.486 | 2.153 | 0.000 | 1.115 | 1.195 | 0.850 | 0.856 | 1.112 | 1.042 | 1.259 | 1.443 | 1.020 | 1.611 | 1.982 | 1.254 | 1.397 | 1.522 | 1.097 |
| C6 | 1.721 | 1.904 | 1.967 | 1.823 | 1.115 | 0.000 | 1.340 | 1.292 | 0.769 | 0.906 | 1.170 | 1.297 | 1.244 | 0.855 | 1.800 | 2.086 | 1.585 | 1.390 | 1.404 | 0.765 |
| C7 | 1.257 | 2.048 | 1.284 | 2.279 | 1.195 | 1.340 | 0.000 | 0.779 | 1.064 | 0.952 | 1.482 | 1.000 | 1.568 | 1.011 | 1.505 | 1.418 | 0.953 | 0.904 | 1.054 | 1.405 |
| C8 | 1.075 | 1.894 | 1.120 | 2.450 | 0.850 | 1.292 | 0.779 | 0.000 | 0.925 | 1.005 | 1.015 | 0.732 | 1.428 | 0.976 | 1.628 | 1.743 | 1.051 | 0.842 | 0.972 | 1.317 |
| C9 | 1.213 | 1.450 | 1.500 | 1.691 | 0.856 | 0.769 | 1.064 | 0.925 | 0.000 | 0.528 | 0.575 | 0.823 | 0.876 | 0.560 | 1.897 | 2.372 | 1.329 | 0.897 | 0.927 | 0.535 |
| C10 | 1.320 | 1.412 | 1.407 | 1.630 | 1.112 | 0.906 | 0.952 | 1.005 | 0.528 | 0.000 | 0.757 | 0.734 | 0.926 | 0.598 | 2.055 | 2.394 | 1.335 | 0.818 | 0.894 | 0.771 |
| C11 | 1.255 | 1.210 | 1.368 | 1.561 | 1.042 | 1.170 | 1.482 | 1.015 | 0.575 | 0.757 | 0.000 | 0.878 | 0.755 | 0.850 | 2.231 | 2.901 | 1.373 | 1.032 | 1.053 | 0.692 |
| C12 | 1.058 | 1.674 | 1.105 | 2.077 | 1.259 | 1.297 | 1.000 | 0.732 | 0.823 | 0.734 | 0.878 | 0.000 | 1.232 | 0.813 | 2.059 | 2.377 | 1.132 | 0.773 | 0.720 | 1.166 |
| C13 | 1.761 | 1.029 | 2.029 | 1.245 | 1.443 | 1.244 | 1.568 | 1.428 | 0.876 | 0.926 | 0.755 | 1.232 | 0.000 | 0.916 | 2.501 | 2.954 | 1.868 | 1.085 | 1.235 | 0.704 |
| C14 | 1.406 | 1.431 | 1.640 | 1.836 | 1.020 | 0.855 | 1.011 | 0.976 | 0.560 | 0.598 | 0.850 | 0.813 | 0.916 | 0.000 | 1.819 | 2.269 | 1.385 | 1.073 | 0.912 | 0.716 |
| C15 | 1.979 | 3.015 | 2.259 | 3.411 | 1.611 | 1.800 | 1.505 | 1.628 | 1.897 | 2.055 | 2.231 | 2.059 | 2.501 | 1.819 | 0.000 | 1.407 | 1.821 | 1.973 | 2.147 | 2.065 |
| C16 | 2.415 | 4.176 | 2.386 | 3.205 | 1.982 | 2.086 | 1.418 | 1.743 | 2.372 | 2.394 | 2.901 | 2.377 | 2.954 | 2.269 | 1.407 | 0.000 | 1.683 | 2.293 | 2.390 | 2.343 |
| C17 | 0.897 | 1.972 | 0.974 | 2.627 | 1.254 | 1.585 | 0.953 | 1.051 | 1.329 | 1.335 | 1.373 | 1.132 | 1.868 | 1.385 | 1.821 | 1.683 | 0.000 | 0.827 | 1.147 | 1.942 |
| C18 | 0.930 | 1.657 | 0.787 | 2.218 | 1.397 | 1.390 | 0.904 | 0.842 | 0.897 | 0.818 | 1.032 | 0.773 | 1.085 | 1.073 | 1.973 | 2.293 | 0.827 | 0.000 | 0.727 | 1.313 |
| C19 | 1.245 | 1.757 | 1.068 | 1.958 | 1.522 | 1.404 | 1.054 | 0.972 | 0.927 | 0.894 | 1.053 | 0.720 | 1.235 | 0.912 | 2.147 | 2.390 | 1.147 | 0.727 | 0.000 | 1.317 |
| C20 | 1.777 | 1.382 | 2.072 | 1.383 | 1.097 | 0.765 | 1.405 | 1.317 | 0.535 | 0.771 | 0.692 | 1.166 | 0.704 | 0.716 | 2.065 | 2.343 | 1.942 | 1.313 | 1.317 | 0.000 |

| **Table S6**. Enrichment values for Glide, Gold and rDock for the selected ensembles (both NMR and X-Ray) structures. | | | |
| --- | --- | --- | --- |
| **Programs** | **Glide** | **Gold** | **rDock** |
| **X-Ray Structures** | | | |
| **16s rRNA A-Site (1)** | | | |
| 2PWT | **24.63** | 22.60 | 26.14 |
| 1YRJ | 20.16 | 26.33 | 24.05 |
| 4F8U | 20.44 | 24.91 | 27.42 |
| Consensus (Mean) | 20.82 | 25.33 | 25.12 |
| Consensus (Best) | 22.10 | 24.67 | *30.13* |
| **PreQ1 Riboswitch** | | | |
| 6E1W | 56.14 | 37.82 | 56.18 |
| 6E1S | 41.74 | 28.99 | 45.08 |
| 7E9E | 56.54 | 39.50 | 58.69 |
| Consensus (Mean) | 46.24 | 36.67 | 52.35 |
| Consensus (Best) | 58.87 | 34.94 | 51.54 |
| **TPP Riboswitch** | | | |
| 2GDI | **47.48** | 31.17 | 45.87 |
| 2HOP | 42.11 | **35.12** | 53.77 |
| 4NYG | 42.89 | 31.45 | 56.18 |
| Consensus (Mean) | 41.07 | 32.29 | 51.08 |
| Consensus (Best) | *47.78* | 32.72 | 49.34 |
| **NMR Conformers** | | | |
| **Tau Pre-mRNA Hairpin** | | | |
| C1 | 14.97 | 16.21 | 31.43 |
| C12 | **20.00** | 17.36 | 31.70 |
| C15 | 13.34 | 15.71 | 29.25 |
| Consensus (Mean) | 16.11 | 16.86 | 33.51 |
| Consensus (Best) | 15.46 | 17.38 | 32.97 |
| **EV71 IRES** | | | |
| C1 | 11.73 | 20.47 | 57.20 |
| C5 | **26.39** | **30.81** | 58.70 |
| C9 | 12.79 | 20.42 | 57.20 |
| Consensus (Mean) | *17.09* | *30.08* | 58.70 |
| Consensus (Best) | *20.34* | *31.18* | 58.70 |
| **HIV FSS** | | | |
| C1 | 30.64 | 23.50 | **23.75** |
| C4 | **33.89** | 24.05 | 19.58 |
| C16 | 28.17 | 29.39 | 18.46 |
| Consensus (Mean) | 32.22 | *30.78* | 19.13 |
| Consensus (Best) | 32.76 | 25.57 | 19.90 |
| The best-performing individual method (10% better than the median) for each target and method is shown in bold; the consensus enrichment method 10% better than the median of the three individual methods is shown in italics; the consensus enrichment method within 10% of the best-performing individual methods are underlined. | | | |

| **Table S7**. Cluster populations from HDBSCAN clustering analysis of RNA binders. | | | | | |
| --- | --- | --- | --- | --- | --- |
| Cluster | Population | In Targets: Intercalating | In Targets: Nucleotide-Like Binding | In Targets: Groove Binding | In Targets: Misc. |
| 1 | 54 | 12 | 0 | 42 | 0 |
| 2 | 29 | 29 | 0 | 0 | 0 |
| 3 | 82 | 0 | 79 | 3 | 0 |
| 4 | 27 | 0 | 0 | 27 | 0 |
| 5 | 27 | 2 | 7 | 12 | 6 |
| 6 | 48 | 34 | 0 | 14 | 0 |
| 7 | 26 | 6 | 0 | 20 | 0 |
| 8 | 32 | 4 | 3 | 25 | 0 |
| 9 | 42 | 10 | 4 | 28 | 0 |
| 10 | 39 | 2 | 0 | 37 | 0 |
| Outliers | 26 | 7 | 11 | 8 | 0 |
